## Supplementary Figures for "Viral modulation of type II interferon increases T cell adhesion and virus spread"

Supplementary Figure 1

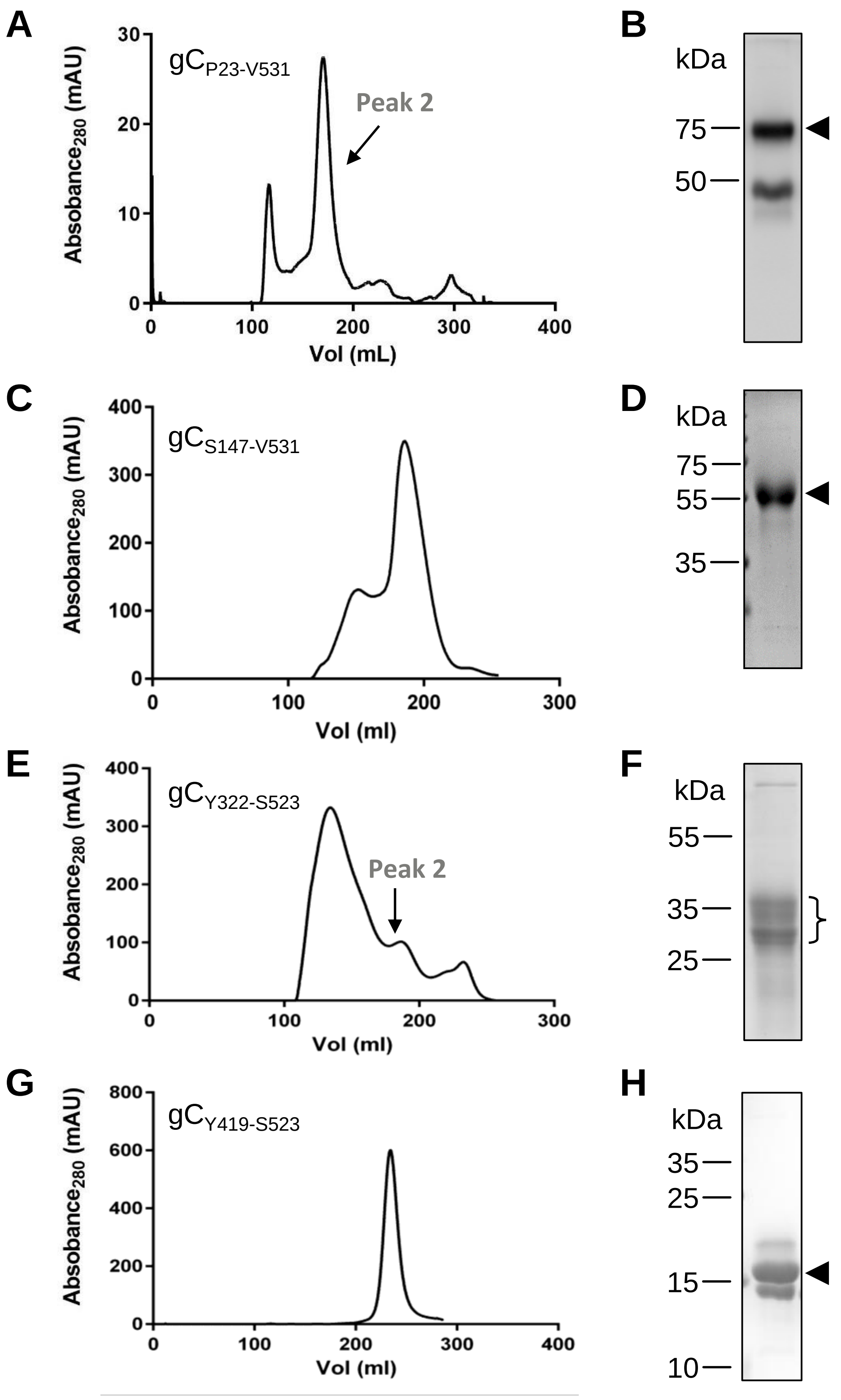

Supplementary Figure 2

A

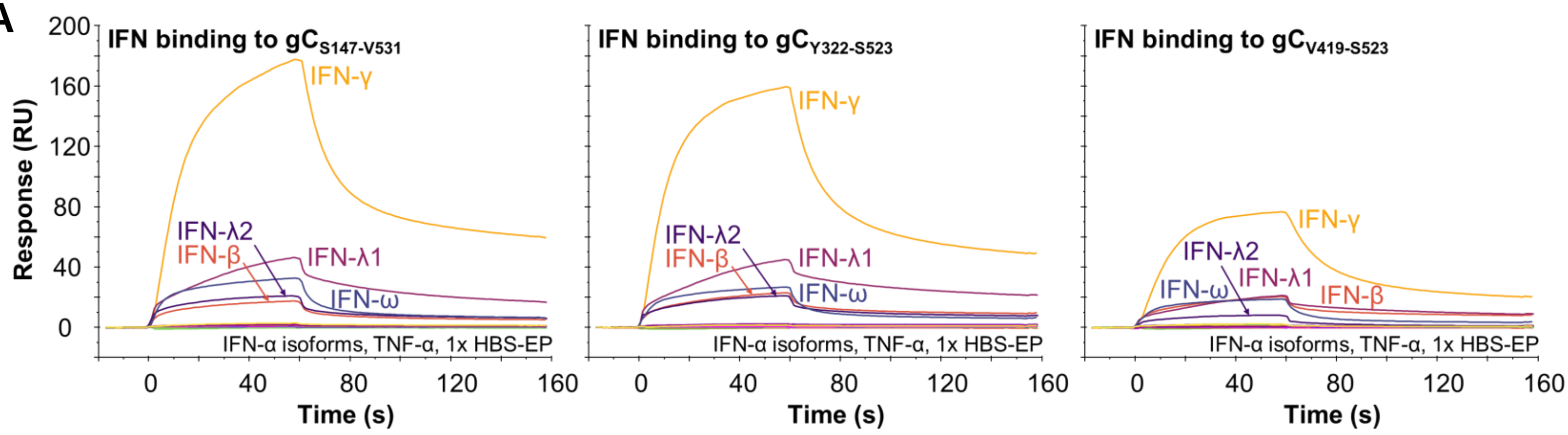


B

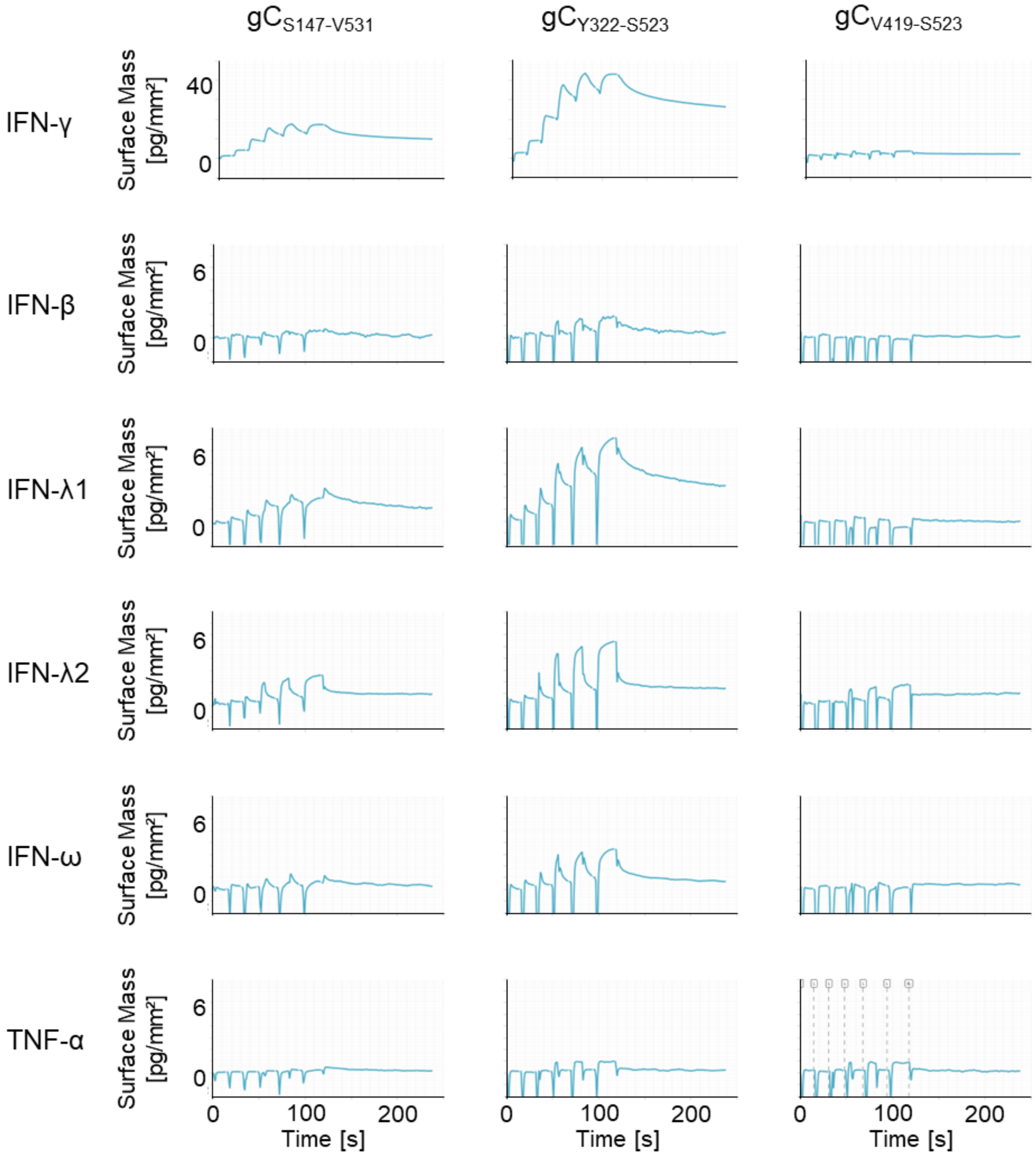

### Supplementary Figure 3

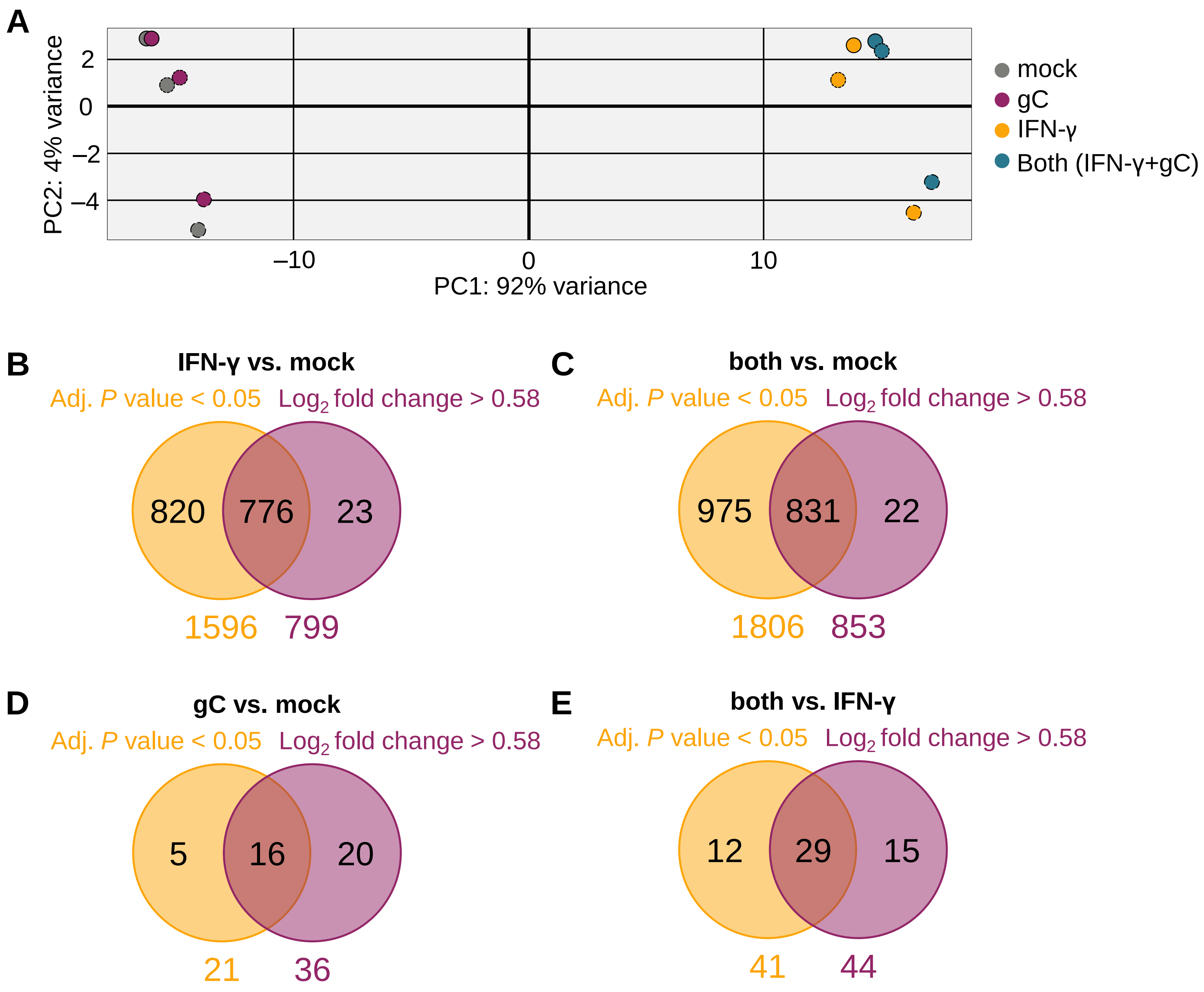

### Supplementary Figure 4

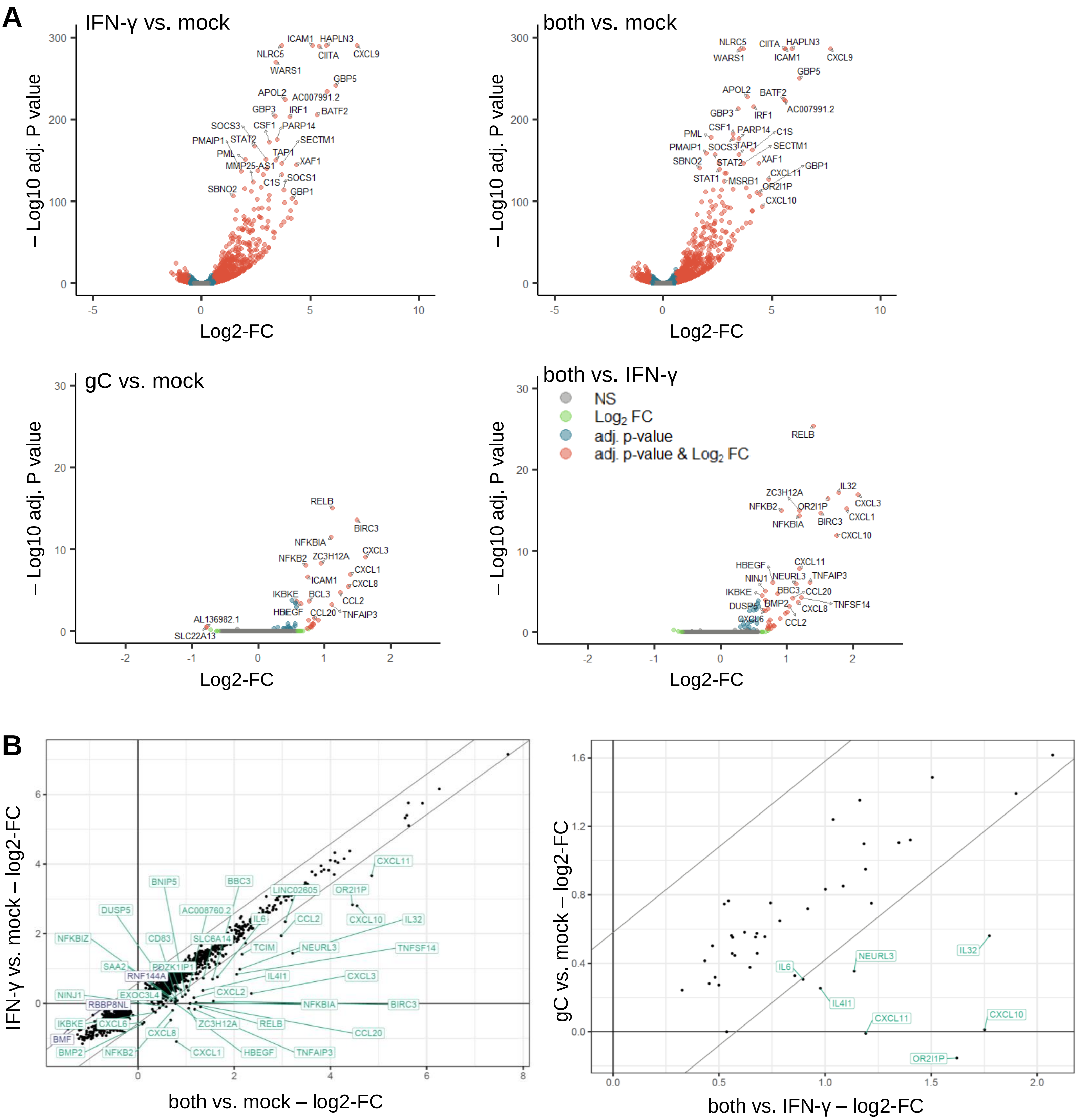

Supplementary Figure 5

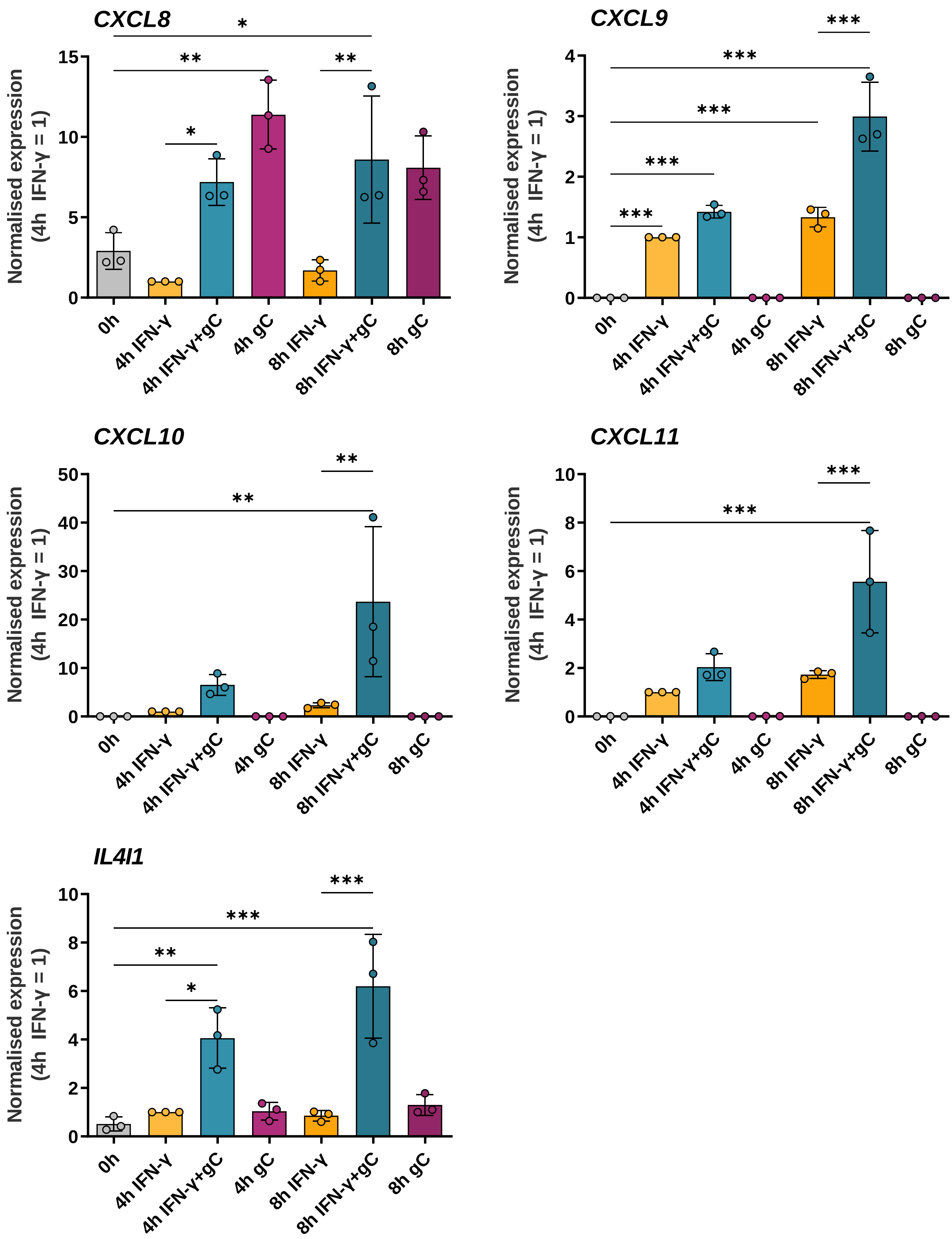

### Supplementary Figure 6

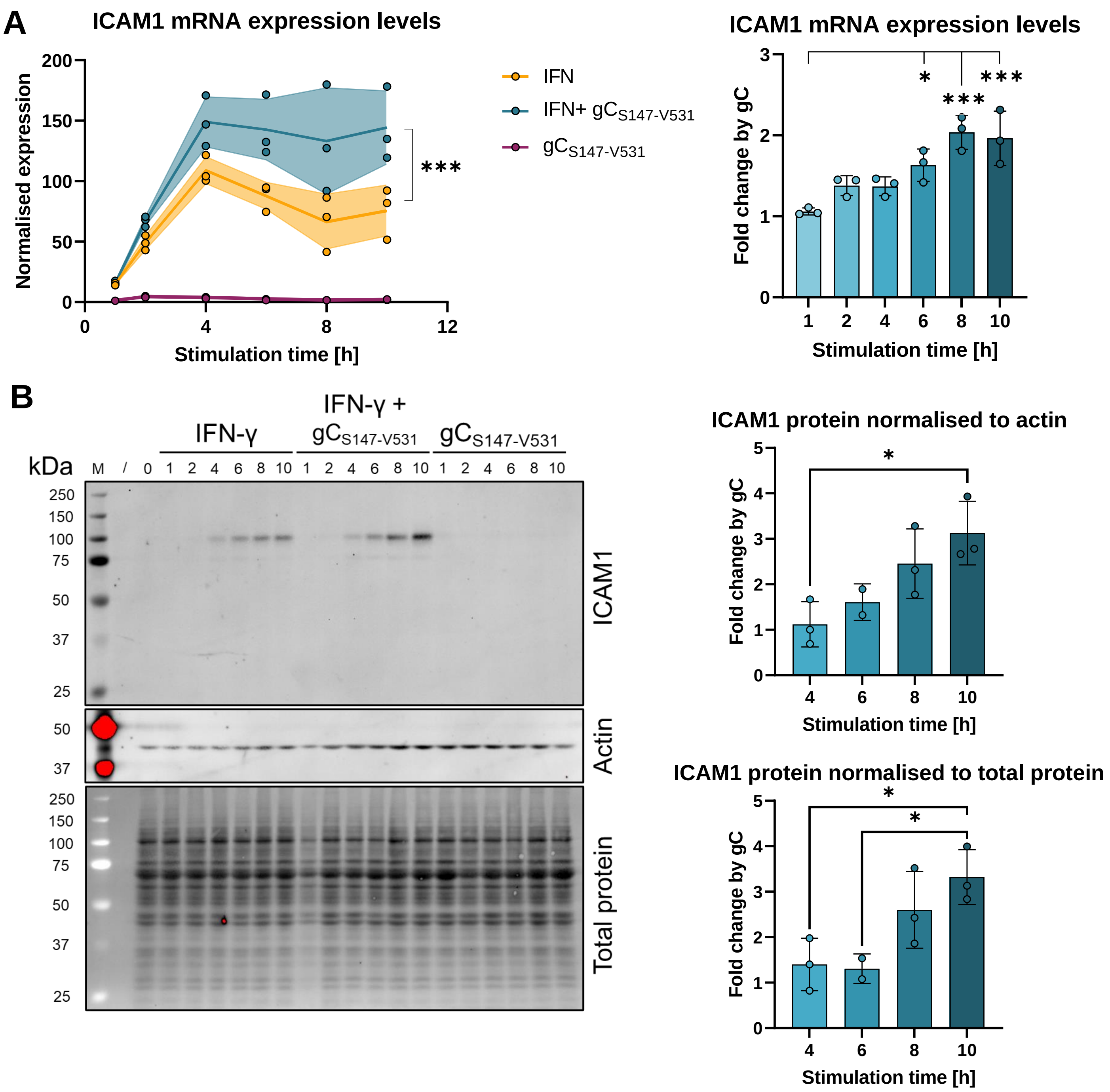

### Supplementary Figure 7

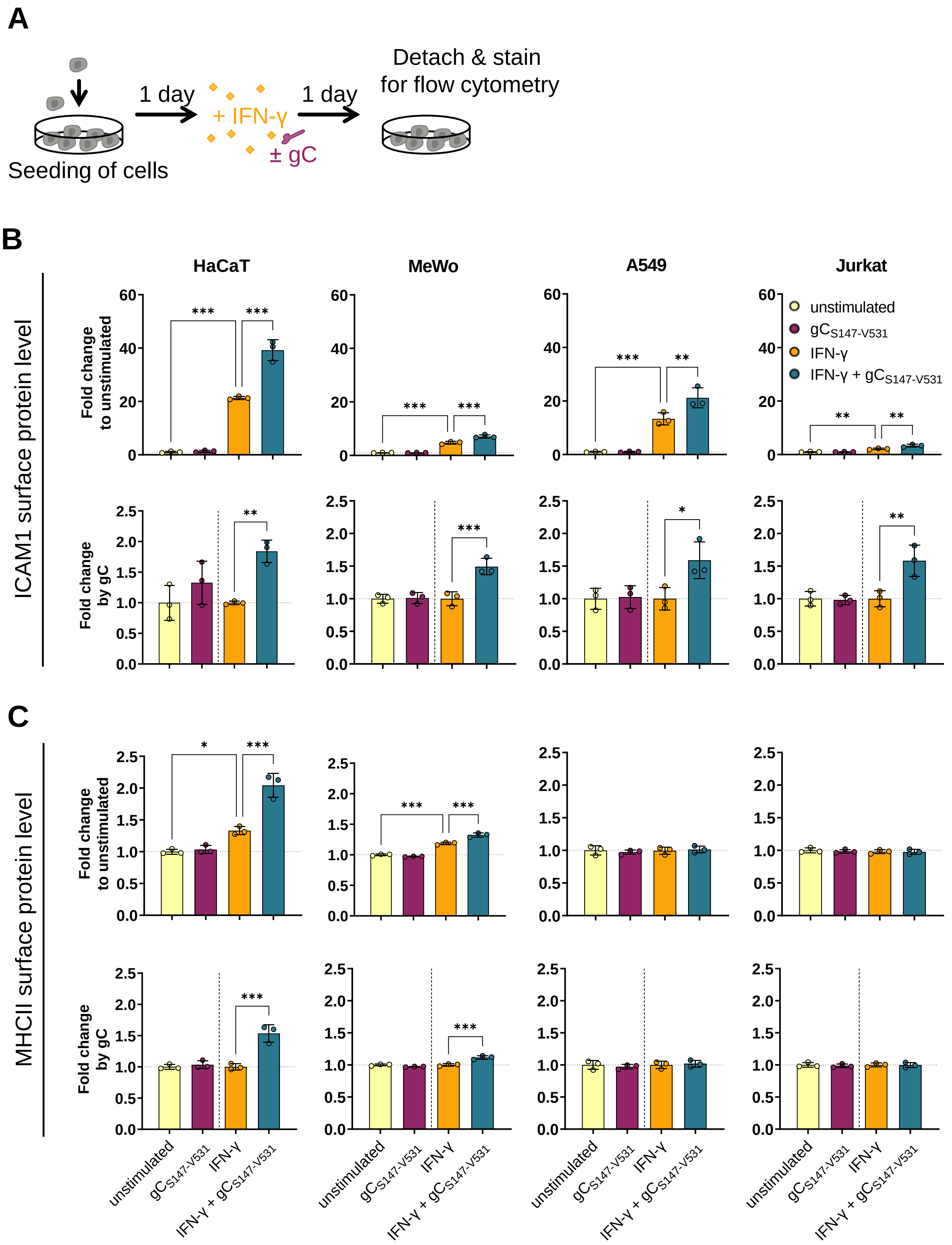

### Supplementary Figure 8

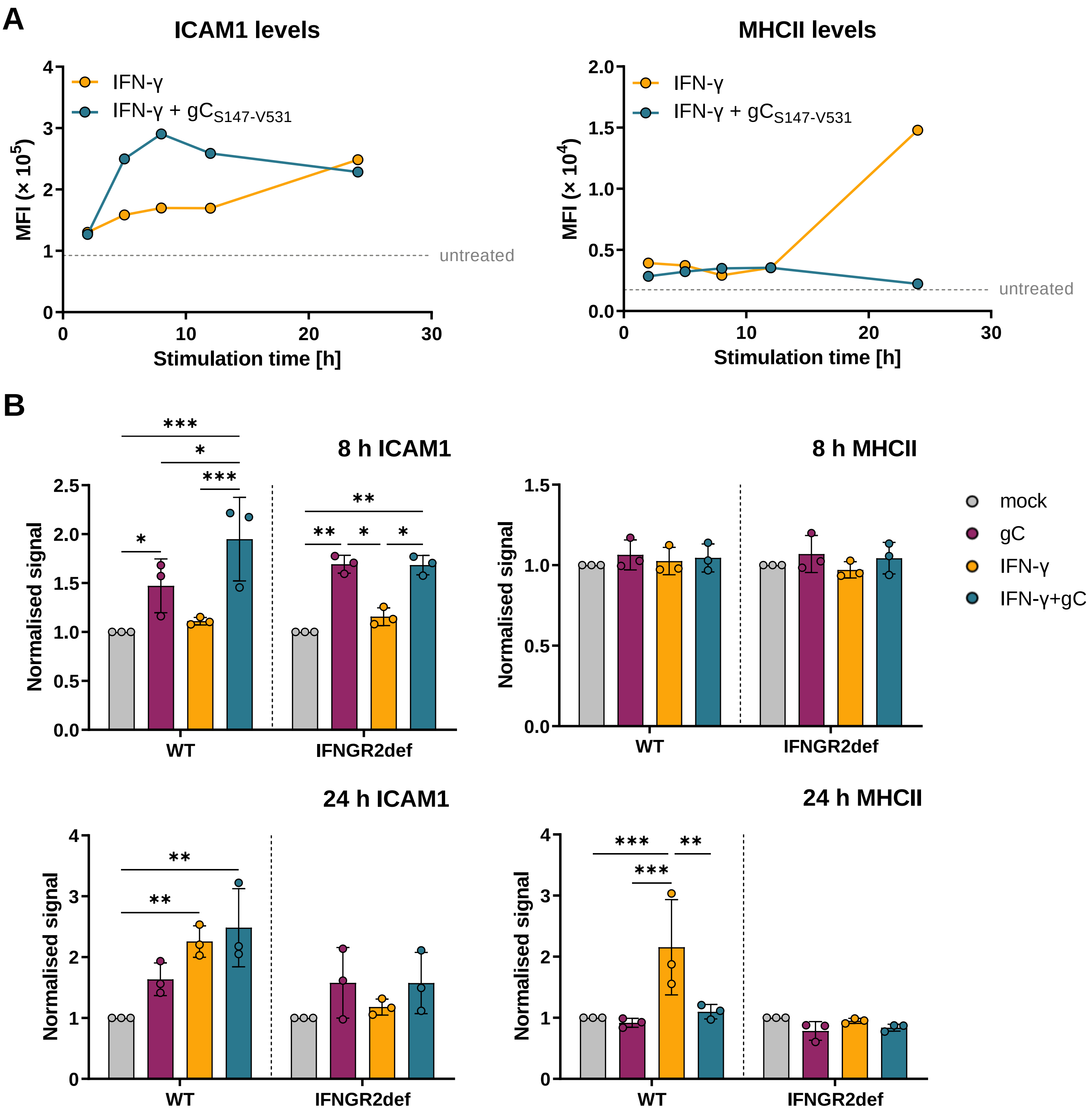

Supplementary Figure 9

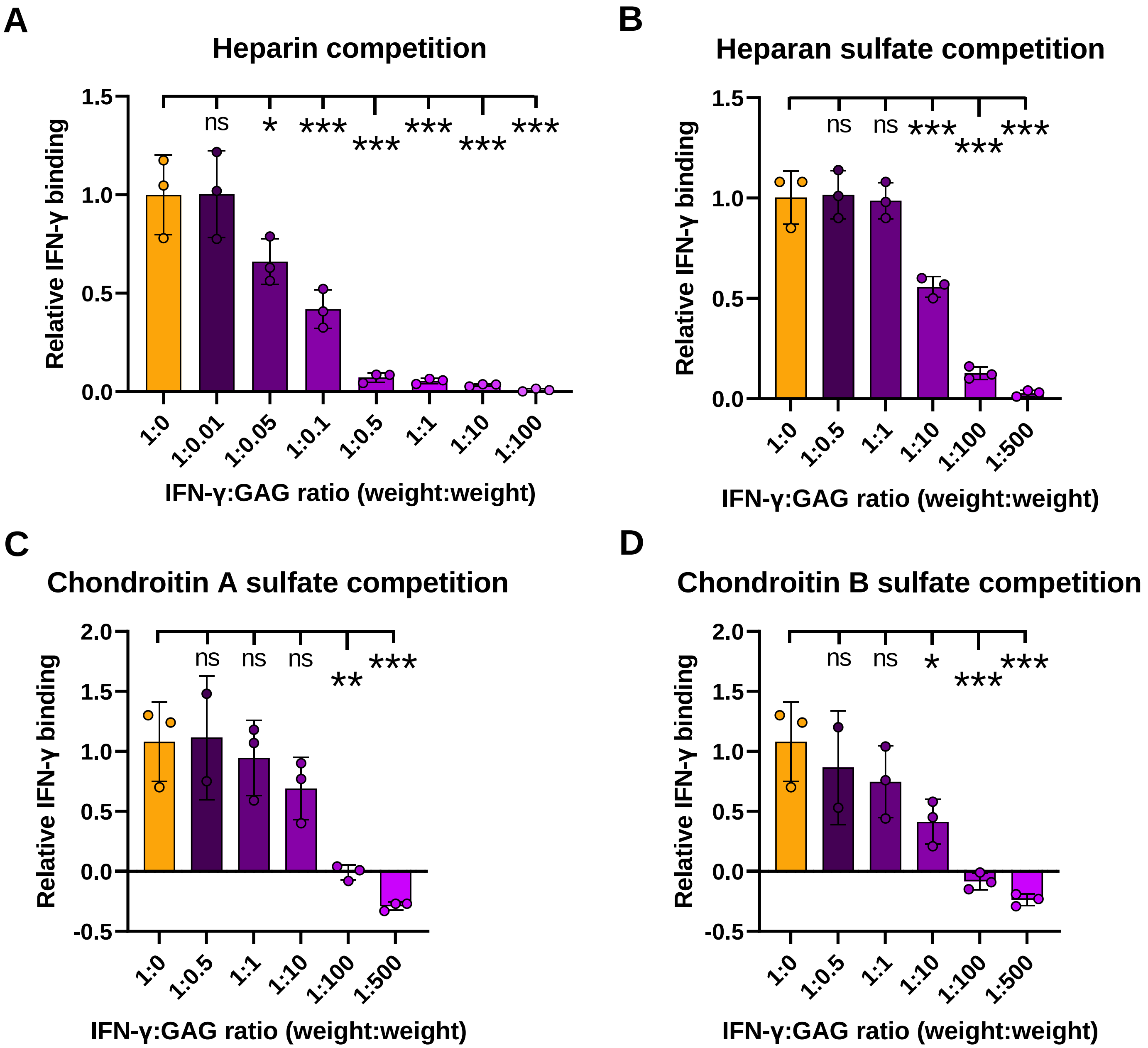

### Supplementary Figure 10

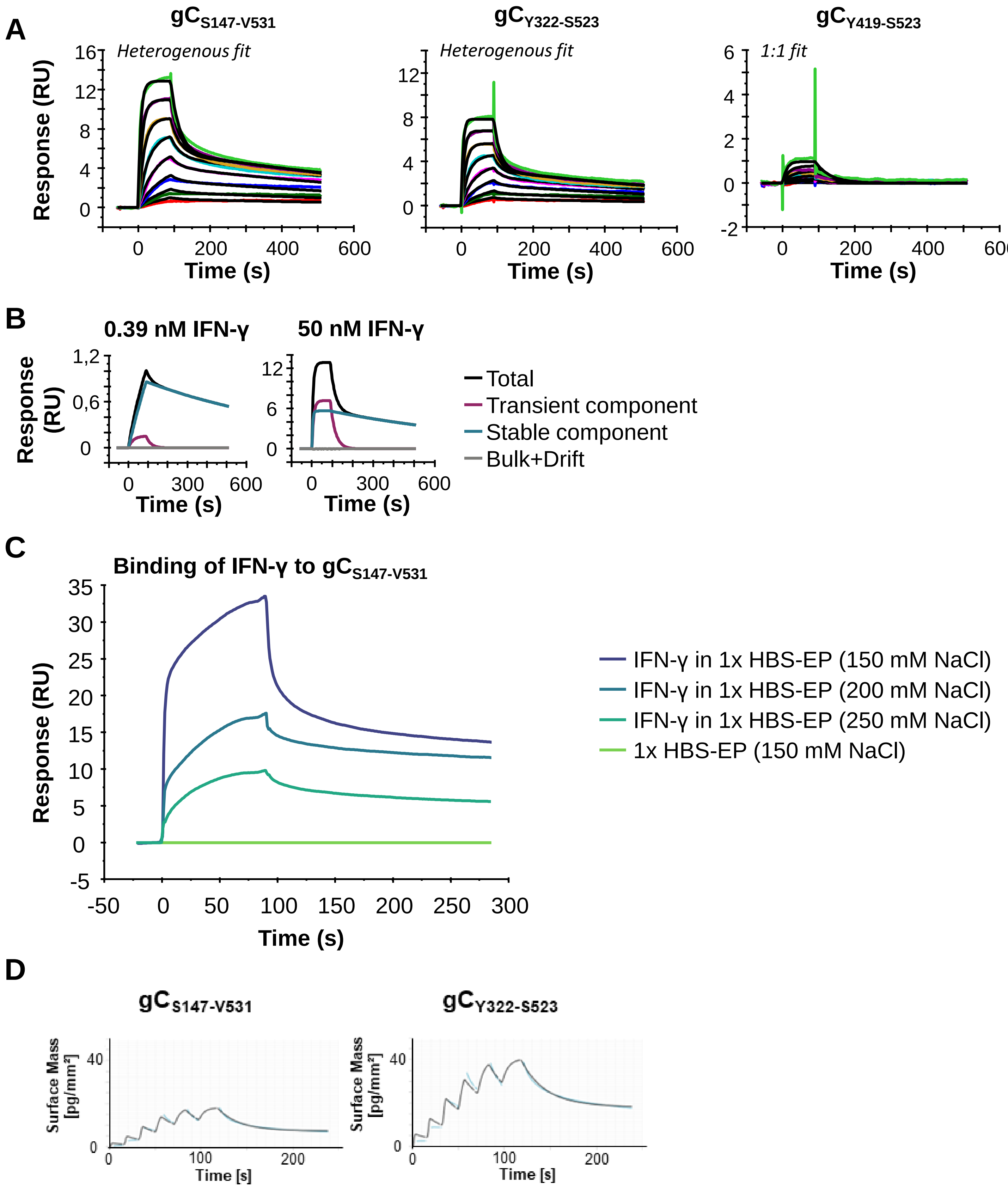

### Supplementary Figure 11

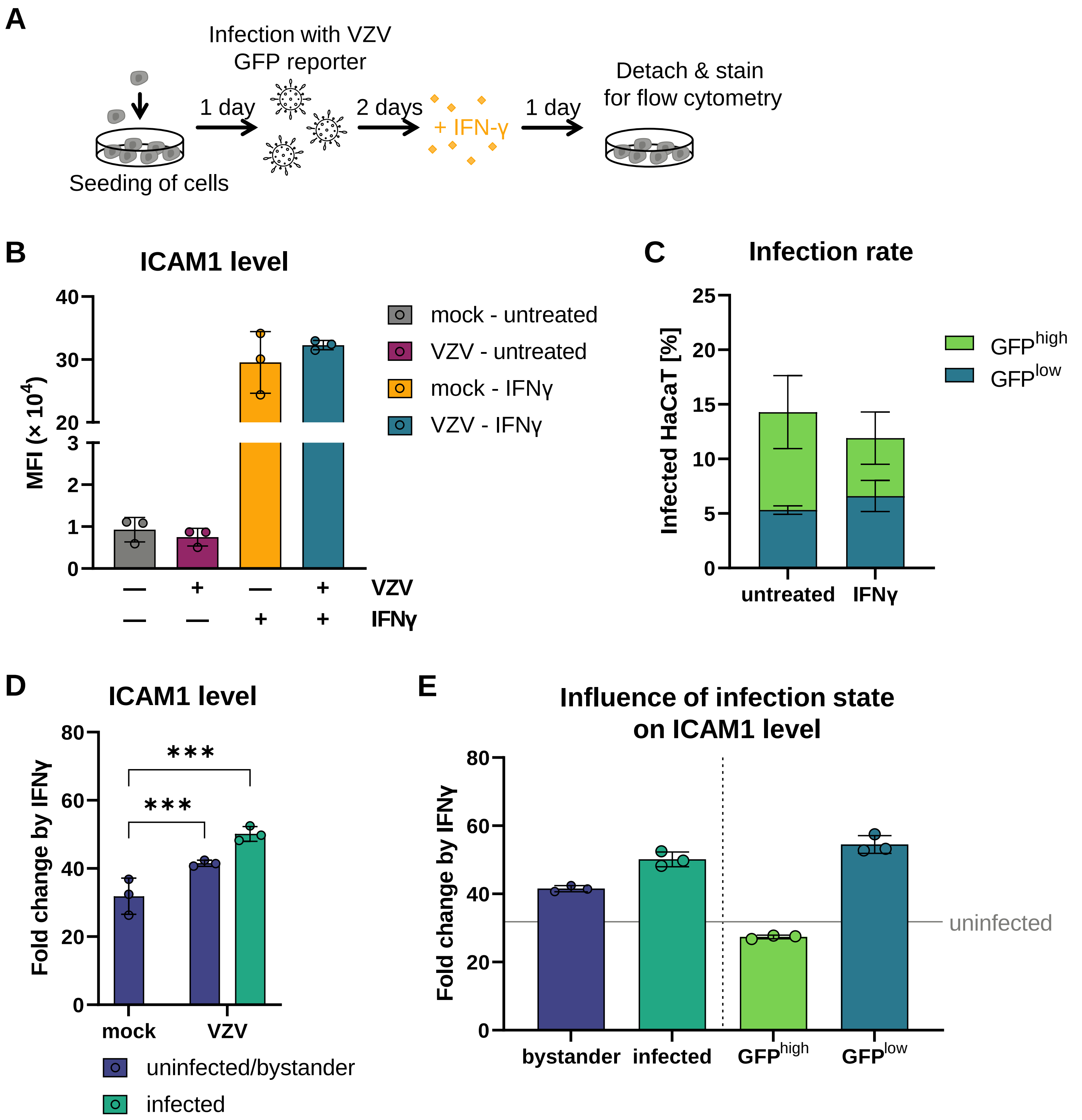

### Supplementary Figure 12

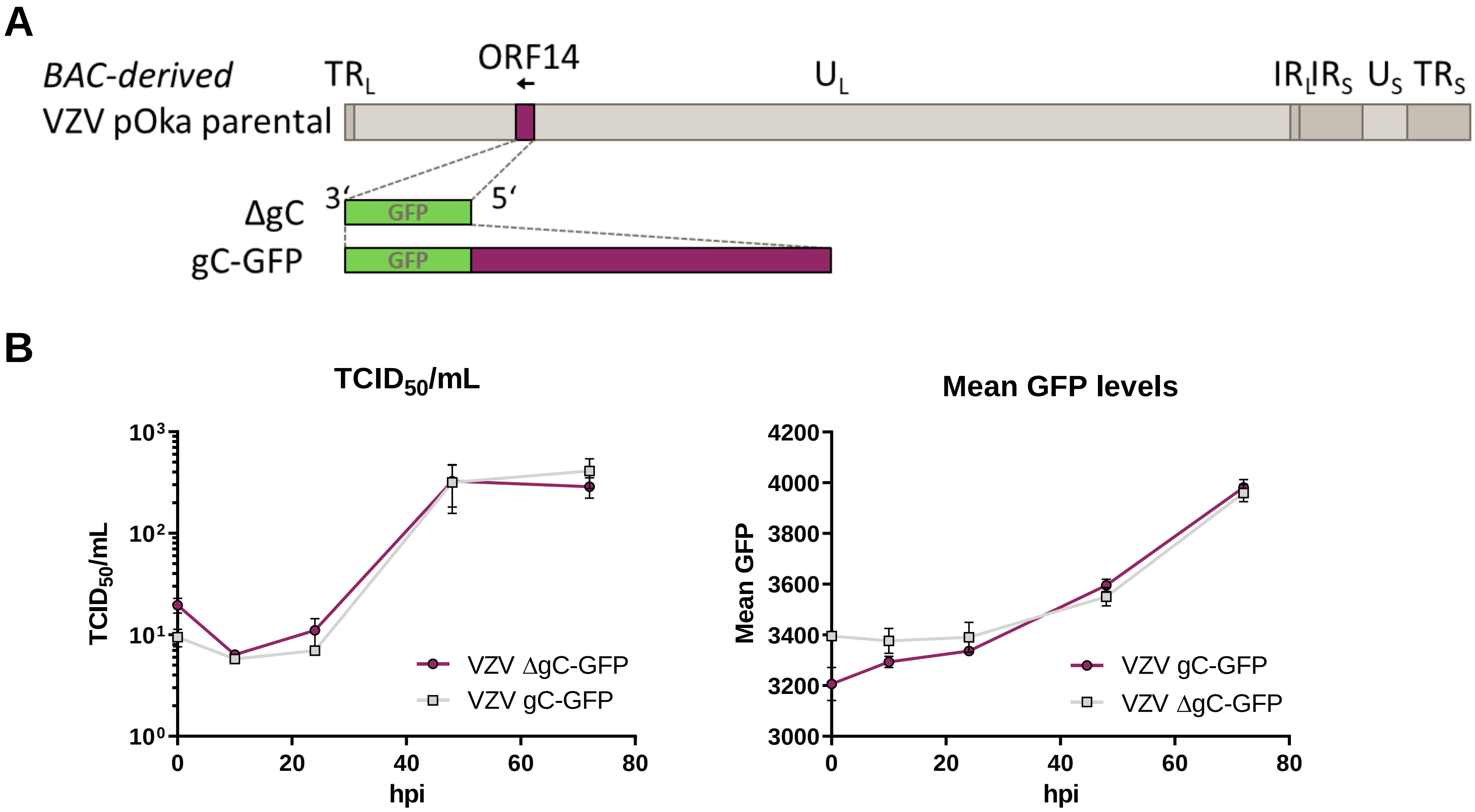
