## Supplementary Tables for "Viral modulation of type II interferon increases T cell adhesion and virus spread"

### Supplementary Table 1

| | Model | Ka1 (1/Ms) | SE(ka1) (1/Ms) | Kd1 (1/s) | SE(kd1) (1/s) | KD1 (M) | Ka2 (1/Ms) | SE(ka2) (1/Ms) | Kd2 (1/s) | SE(kd2) (1/s) | KD2 (M) |
|---|---|---|---|---|---|---|---|---|---|---|---|
| **$gC_{S147\text{-}V531}$** | Heterogeneous Ligand | 1,51E+06 | 7,76E+03 | 4,14E-02 | 1,39E-04 | 2,75E-08 | 4,93E+06 | 7,14E+03 | 1,11E-03 | 3,90E-06 | 2,26E-10 |
| **$gC_{Y322\text{-}S523}$** | Heterogeneous Ligand | 2,61E+06 | 1,50E+04 | 5,58E-02 | 2,35E-04 | 2,13E-08 | 6,47E+06 | 1,29E+04 | 1,33E-03 | 4,89E-06 | 2,05E-10 |
| **$gC_{Y419\text{-}S523}$** | 1:1 Binding | 1,35E+06 | 1,88E+04 | 2,44E-02 | 1,77E-04 | 1,80E-08 | n.a. | n.a. | n.a. | n.a. | n.a. |

Suppl. Table 1
Table showing exemplarily kinetic values for the VZV gC – IFN-γ interaction obtained from at least two kinetic assays performed using the Biacore S200. Abbreviations: s = seconds, M = molar (mol/L), Ka = association rate constant, Kd = dissociation rate constant, KD = equilibrium dissociation constant, SE = standard error.

### Supplementary Table 2

| | Model | Ka1 (1/Ms) | Kd1 (1/s) | KD1 (M) | Ka2 (1/Ms) | Kd2 (1/s) | KD2 (M) |
|---|---|---|---|---|---|---|---|
| **$gC_{S147\text{-}V531}$** | Heterogeneous Ligand | 7,17E+05 | 4,29E-02 | 5,98E-08 | 8,06E+05 | 1,57E-05 | 1,95E-11 |
| **$gC_{Y322\text{-}S523}$** | Heterogeneous Ligand | 6,18E+05 | 3,91E-02 | 6,33E-08 | 6,84E+05 | 3,28E-04 | 4,80E-10 |

Suppl. Table 2
Table showing exemplarily kinetic values for the VZV gC – IFN-γ interaction obtained from at three kinetic assays performed using the Creoptix WAVE systems. Abbreviations: s = seconds, M = molar (mol/L), Ka = association rate constant, Kd = dissociation rate constant, KD = equilibrium dissociation constant.
