## Supplementary Figure Legends for "Viral modulation of type II interferon increases T cell adhesion and virus spread"

**Supplementary Figure 1. Purification of recombinant VZV gC proteins expressed in *Drosophila* S2 cells. (A, C, E, G)** Size-exclusion chromatography (SEC) profiles of purified recombinant gC$_{P23\text{-}V531}$ (A), gC$_{S147\text{-}V531}$ (C), gC$_{Y322\text{-}S523}$ (E) and gC$_{Y419\text{-}S523}$ (G) expressed in *Drosophila* S2 cells. Purification details can be found in Materials and Methods. **(B, D, F, H)** Images taken from SDS-PAGE loaded with purified gC$_{P23\text{-}V531}$ (B), gC$_{S147\text{-}V531}$ (D), gC$_{Y322\text{-}S523}$ (F) and gC$_{Y419\text{-}S523}$ (H) and stained for total protein with TCE (B) or Coomassie (D, F, G). In (B) and (F) the analyzed sample was obtained from peak 2. The molecular weight marker (in kDa) is shown on the left side.

**Supplementary Figure 2. gC binds IFN-γ better than the other IFNs. (A, B)** Sensorgrams showing the results of binding screenings between VZV gC constructs and cytokines using the Biacore S200 (A) or the Creoptix WAVE (B) systems. In (A) the gC constructs were immobilized on a CM5 sensor chip (2853 RU, 1623 RU, and 668 RU, respectively) and the cytokines were injected at 100 nM with a flow rate of 30 µL/min. In (B) the different gC proteins were immobilised on DXH chips (567 pg/mm$^3$, 584 pg/mm$^3$, and 296 pg/mm$^3$, respectively) and cytokines were injected at 200 nM using RAPID in the tight binder mode. Please note the different Y-axis scaling for IFN-γ compared to the other cytokines. Abbreviations: s = seconds, RU = resonance units

**Supplementary Figure 3. VZV gC modifies the expression of a low number of genes, including some modulated by IFN-γ.** HaCaT cells were stimulated with IFN-γ, gC$_{S147\text{-}V531}$, both or mock treated for 4 h. RNA was isolated and further processed for RNAseq. **(A)** Principal component analysis (PCA) of gene counts of stimulated HaCaT cells from three biological replicates. IFN-γ had the strongest effect on variation in the samples, whereas gC only had minor effects on gene expression compared to mock. Similar boundaries of circles indicate samples obtained from the same experiment. **(B-E)** Venn diagrams showing the number of genes, whose expression level was modified at least 1.5-fold (purple) in a statistically significant manner (*P* value < 0.05; yellow) between the two compared conditions. Differential gene expression analysis was performed comparing the different treatment conditions.

**Supplementary Figure 4. VZV gC induces ISG expression in a biased manner.** HaCaT cells were stimulated with IFN-γ, $gC_{S147\text{-}V531}$, both or mock treated for 4 h. RNA was isolated and further processed for RNAseq. Differential gene expression analysis was performed using DESeq2 comparing the different treatment conditions as indicated. **(A)** Volcano plots; x-axis represents the log2 of the fold change (FC), y-axis represents the negative decade logarithm of the adjusted (adj.) *P* value for the four different comparisons. Each circle represents a gene. Boundaries were set with adj. *P* value of 0.05 and a fold change of 1.5 (equals log2-FC of 0.58). Red circles represent genes with significant change in the adj. *P* value and the fold change, blue circles depict genes with only a significant adj. *P* value, and green circles represent genes with a significant change only in the fold change. In the representation of the IFN-γ-treated versus mock control comparison, the adj. *P* values for *CXCL9*, *HAPLN3*, *NLRC5* and *ICAM1* were too low to be calculated, hence they were set to the lowest calculated adj. *P* value (*CIITA*) in this data set. In the representation of the comparison of HaCaT cells treated with IFN-γ and gC versus mock-treated cells, the adj. *P* values for *CXCL9*, *HAPLN3*, *CIITA*, *NLRC5* and *ICAM1* were too low to be calculated, hence they were set to the lowest calculated adj. *P* value (*WARS1*) in this data set. **(B)** Graphs showing the fold changes (FC) of significantly regulated genes from the indicated comparisons. The grey lines indicate a corridor in which genes are less than 1.5-fold changed between the two groups. Genes below the corridor are more upregulated in the comparison plotted at the x-axis (green). Genes above the corridor are stronger regulated in the comparison plotted at the y-axis (violet).

**Supplementary Figure 5. Validation of the RNASeq data by RT-qPCR.** Graphs showing expression of *CXCL8*, *CXCL9*, *CXCL10*, *CXCL11* and *IL4I1* relative to *actin* in HaCaT cells stimulated with 5 ng/mL IFN-γ, 300 nM gC$_{S147-V531}$ or both for the indicated time points. The values were normalized to those obtained from cells stimulated for 4 h with IFN-γ. Each filled circle corresponds to one independent assay. Error bars represent standard deviation of the arithmetic mean from three independent experiments. One-way ANOVA was performed, followed by Šídák's multiple comparisons (comparing each sample to the mock and between IFN-γ stimulated cells vs. co-stimulated cells for each time point). Non-significant comparisons are not indicated. * = $P$ <0.033; ** = $P$ <0.002; *** = $P$ <0.001.

**Supplementary Figure 6. HaCaT cells co-stimulated with VZV gC and IFN-γ show increased mRNA expression and total ICAM1 protein levels. (A)** HaCaT cells were stimulated with 5 ng/mL IFN-γ and/or 300 nM gC$_{S147-V531}$ for the indicated time points, RNA was isolated and analyzed for *ICAM1* expression by RT-qPCR. Using the ΔΔCt method, *ICAM1* level was normalised to unstimulated cells and to actin as housekeeping gene. Shown are the individual values from three independent assays (filled circles), the resulting mean is shown as solid line with standard deviation (SD) as colored transparent background. The area under the curve (AUC) was calculated, then one-way ANOVA was performed on the AUC values, followed by Dunnett's multiple comparisons. The fold change induced by addition of gC$_{S147-V531}$ to IFN-γ was calculated for each timepoint and plotted in a bar chart (right panel). Filled circles represent the individual values from each independent experiment, bars represent the mean ± SD. One-way ANOVA, followed by Dunnett's multiple comparisons was performed (comparing to the 1 h stimulation time). **(B)** Immunoblots detecting ICAM1 (top panel) and actin (middle panel) and blot showing total protein signal visualised using trichlorethanol (bottom panel) in same samples employed in (A). Quantification of the western blot band intensities is shown in the right panels. Top panel shows the normalization of ICAM1 signal to actin, bottom panel shows ICAM1 signal normalized to total protein. One representative experiment out of three biological repeats is shown. One-way ANOVA, followed by Tukey's multiple comparisons was performed (comparing all samples to each other). ns = not significant; * = *P* <0.033; ** = *P* <0.002; *** = *P* <0.001. Abbreviations: kDa = kilo Dalton; M = marker; h = hours

**Supplementary Figure 7. Cell lines co-stimulated with VZV gC and IFN-γ have increased surface levels of ICAM1. (A)** Schematic representation of the assay. Cells were seeded one day prior to stimulation with 5 ng/mL IFN-γ, 300 nM VZV $gC_{S147-V531}$ or both. 24 h after stimulation the cells were detached, stained, and analyzed by flow cytometry to detect ICAM1 (B) of MHCII (C) at the plasma membrane. **(B, C)**. Bar charts showing the fold change of ICAM1 (B) or MHCII (C) surface protein levels compared to unstimulated cells (top row) or induced by $gC_{S147-V531}$ compared to either mock or IFN-γ baseline (bottom row). The median fluorescence intensities were determined after gating on single and alive cells. Bars show the mean ± SD, filled circles represent values from three independent experiments. One-way ANOVA, followed by Šídák's multiple comparisons was performed (comparison IFN-γ to unstimulated and/or condition with gC to baseline without gC). Non-significant comparisons are not indicated. * = *P* <0.033; ** = *P* <0.002; *** = *P* <0.001.

**Supplementary Figure 8. Effect of gC on ICAM1 protein level iPSC-derived macrophages. (A, B)** iPSC-derived macrophages from a healthy donor (A, B) and from and IFNGR2-deficient donor (B) were mock-stimulated or stimulated with 5 ng/mL IFN-γ, 300 nM gC$_{S147-V531}$ or both for the indicated time points and then labelled with antibodies to ICAM1, MHCII, and stained with Zombie-NIR dye. **(A)** Graphs showing ICAM1 (left) and MHCII (right) protein levels on the plasma membrane. Cells were analyzed by flow cytometry and median fluorescence intensities (MFI) were determined after gating on alive single cells. **(B)** Graphs showing ICAM1 (left) and MHCII (right) protein levels at the plasma membrane at 8 and 24 hours post-stimulation. MFI was normalized to the mock-treated cells for each donor within each experiment. Circles represent results of biological replicates, bars show mean values ± SD. Two-way ANOVA, followed by Tukey’s multiple comparisons was performed (comparison between all treatments within each donor). Non-significant comparisons are not depicted. * = $P$ <0.033; ** = $P$ <0.002; *** = $P$ <0.001.

**Supplementary Figure 9. GAGs compete with the gC – IFN-γ interaction. (A-D)** Bar graphs showing the effect of heparin (A), heparan sulphate (B), chondroitin A sulphate (C) and chondroitin B sulphate (D) on binding of IFN-γ to gC. Purified gC$_{S147-V531}$ was immobilized on a CM5 sensor chip (9,800 RU) and IFN-γ was injected at 100 nM either alone or together with increasing amounts of different GAGs (weight ratio). The response levels at the binding report point were normalized to the response obtained for IFN-γ alone and plotted for the different weight ratios. The bars represent the mean ± SD, the filled circles represent the values from three independent experiments. One-way ANOVA was performed to test for statistical significance (comparing to control without GAGs), followed by Dunnett’s multiple comparisons test. ns = not significant; * = $P$ <0.033; ** = $P$ <0.002; *** = $P$ <0.001

**Supplementary Figure 10. The interaction between gC and IFN-γ contains a stable and a transient component**

**(A)** Sensorgrams showing the results of assays to determine the kinetics of the interactions between VZV gC constructs and IFN-γ using the Biacore S200. The gC constructs were immobilized on a CM4 sensor chip (877 RU, 399 RU, and 293 RU, respectively) and IFN-γ was injected in a 1:2 dilution series starting at 50 nM with a flow rate of 30 µL/min. Black lines indicate the heterogenous ligand fit or 1:1 binding model (as indicated) determined with the Biacore S200 Evaluation software. **(B)** Sensorgrams showing the contribution of the transient and stable interactions upon injection of a low and high IFN-γ concentration onto the $gC_{S147-V531}$ chip. **(C)** Sensorgram showing results of binding experiments between $gC_{S147-V531}$ immobilised on a CM5 sensor chip (877 RU) and IFN-γ. The A-B-A injection scheme of the Biacore S200 system to test different buffer conditions was used. Pre-sample contact time was set to 360 s. The association time was 90 s, followed by 240 s of post-sample contact time. IFN-γ was injected at 100 nM with a flow rate of 30 µL/min. **(D)** Sensorgrams showing the results of assays to determine the kinetics of the interactions between VZV $gC_{S147-V531}$ and $gC_{Y322-V523}$ using the Creoptix WAVE system. The gC proteins were immobilised on a DXH chip (567 pg/mm$^3$ and 584 pg/mm$^3$, respectively) and IFN-γ was injected at 200 nM using RAPID in the tight binder mode. Black line indicates the fit using a heterogenous ligand model and traditional fitting.

**Supplementary Figure 11. Induction of ICAM1 by IFN-γ is higher in VZV inoculated cultures. (A)** Schematic representation of the assay. HaCaT cells were seeded 24 h prior to infection with pOka-Δ57-GFP. 48 h after infection, the cells were stimulated with IFN-γ or mock treated. The next day, cells were detached and labelled with anti-ICAM1-APC and stained with Zombie-NIR dye, fixed, and analyzed by flow cytometry. **(B)** Bar chart showing the median fluorescence intensities (MFI) for ICAM1 in the four treatment conditions after gating on alive cells. **(C)** Bar chart showing the percentage of infected HaCaT cells in mock- and IFN-γ-treated cultures after gating on alive cells. The different colors discriminate the proportion of $GFP^{high}$ and $GFP^{low}$ expressing cells. Ordinary two-way ANOVA analysis showed no significant differences. **(D,E)** Bar charts showing the calculated fold-change of ICAM1 levels in mock and VZV-infected cells after gating on uninfected or infected cells (D) and after differentiating between $GFP^{high}$ and $GFP^{low}$ cells (E). As reference in E the fold change by IFN-γ of uninfected cells (from D) is indicated with a grey line. Ordinary two-way ANOVA analysis with main effects only followed by Dunett's multiple comparison was performed (D). Bars in (B-E) represent the mean ± SD and filled circles represent the individual values from three independent experiments. ns = not significant; * = $P$ <0.033; ** = $P$ <0.002; *** = $P$ <0.001.

**Supplementary Figure 12. VZV-gC-GFP and VZV-ΔgC-GFP show similar replication kinetics in HaCaT cells. (A)** Schematic representation of the BAC-derived VZV pOka strains. The repeat and unique regions of the parental, BAC-derived, VZV genome are shown. *ORF14*, encoding gC is highlighted in magenta. The arrow above *ORF14* indicates that this gene is located in the reverse DNA strand. Monomeric GFP was inserted instead of the *ORF14* locus and at the 3' end of *ORF14* to generate VZV-ΔgC-GFP and VZV-gC-GFP recombinant viruses, respectively. In both cases, the *ORF14* promoter drives expression of GFP. **(B)** HaCaT cells were infected with 100 PFU of the two indicated BAC-derived virus strains. At the different time points post-infection, cells were imaged and collected for titration. Assays were performed in triplicates. Collected cells were titrated and the $TCID_{50}$ was determined on HaCaT cells. $TCID_{50}$ values are plotted over time in the left panel. The right panel shows the mean GFP fluorescence over time measured with the Cytation3 (BioTek). Abbreviations: $TR_l$ = terminal repeat long; $TR_s$ = terminal repeat short; $U_L$ = unique long region; $IR_L$ = internal repeat long; $IR_s$ = internal repeat short; $U_S$ = unique short region; ORF = open reading frame; GFP = green fluorescent protein; $TCID_{50}$ = tissue culture infectious dose 50; hpi = hours post-infection.
