## Supplementary Materials and Methods for "Viral modulation of type II interferon increases T cell adhesion and virus spread"

**Bacteria**

*Escherichia coli* (*E. coli*) DH5α was grown in LB medium shaking at 200 – 220 rpm or on LB agar at 37 °C. This strain was used for general cloning purposes. The VZV BAC-containing *E. coli* strain GS1783 was cultured in LB medium containing 17 µg/mL chloramphenicol shaking at 200-220 rpm or on LB agar containing 17 µg/mL chloramphenicol at 30-32 °C.

**Cell culture**

A549 is an epithelial cell line isolated from lung tissue of a caucasian male with lung cancer and was provided by Thomas Pietschmann (Twincore, Hannover, Germany). HaCaT cells are *in vitro* spontaneously transformed keratinocytes derived from histologically normal skin from a 1aucasian male. These cells were provided by Beate Sodeik (Institute of Virology, MHH, Germany). Jurkat E6.1 originate from a male patient with acute T cell leukemia and were kindly provided by Martin Messerle (Institute of Virology, MHH, Germany). Jurkat LFA-1 KO cells were a gift from Carsten Münk (Düsseldorf University Hospital, Germany) and have been described previously[1]. MeWo cells are derived from a male patient suffering malignant melanoma and were purchased from ATCC (HTB-65™). Peripheral blood mononuclear cells (PBMCs) from anonymised healthy blood donors were isolated using standard Ficoll-density centrifugation methods. Isolated PBMCs were washed twice, and remaining erythrocytes were lysed with ACK lysing buffer (Lonza). The PBMCs were frozen in 90% FBS and 10% DMSO and freshly thawed before experiments.

All mammalian cells were cultured at 37 °C with 5% $CO_2$ in a humidified incubator. A549, HaCaT and Mewo cells were cultured in DMEM (Gibco™ #41966-052), supplemented with 8% heat-inactivated FBS (Sigma #F7524), 1× L-glutamine (Cytogen #04-80100) and 1× penicillin/streptomycin (Cytogen #06-07100). Jurkat E6.1

and Jurkat LFA-KO cells were cultured in RPMI1640 (Gibco™ #21875-034), supplemented with 8% heat-inactivated FBS, 1× L-glutamine and 1× penicillin/streptomycin, with the addition of 2 µg/mL puromycin (Invivogen #ant-pr) for the Jurkat LFA-KO cells. PBMCs were maintained in RPMI1640 supplemented with 10% heat-inactivated FBS, 1× L-glutamine, 1× sodium pyruvate (Gibco™ #11360-070) and 1× penicillin/streptomycin.

Schneider's *Drosophila melanogaster* Line 2 (S2) cells (ATCC No.: CRL-1963) are derived from embryonic tissue of the fruit fly *Drosophila melanogaster* and were used for protein production after stable transfection. The cell line was purchased from Thermo Scientific. These cells were grown in Schneider's *Drosophila* medium (Gibco) supplemented with heat-inactivated 10% FBS and 1× penicillin/streptomycin (Gibco) at 28 °C with normal atmospheric conditions in a semi-adherent manner for maintenance and transfection. After transfection, 8 µg/mL puromycin was added to the media for selection during maintenance. For protein production, this semi-adherent cell line was grown in suspension by shaking at 70 rpm in baffled Erlenmeyer flasks using Insect XPRESS medium (Lonza).

For the generation of healthy iPSC-derived macrophages, both hCD34iPSC16 (MHHi015-A: https://hpscreg.eu/cell-line/MHHi015-A) (for the experiments depicted in the Supplementary Figure 10 A) or the GMP research grade LiPSC-GR1.1 iPSC line (GMP-grade, Lonza, Basel, Switzerland) (for the experiments depicted in the Supplementary Figure 10B) were used. The patient-specific iPSC line derived from a patient harbouring the *IFNGR2* c.705C>A mutation (citation: 10.3390/cells9020483) was used for the generation of the IFNGR2-deficient macrophages. iPSC-derived macrophages were generated as described previously[2,3]. Briefly, upon expansion of the iPSC cells, bFGF was omitted from the culture medium and embryoid bodies (Ebs) were formed on an orbital shaker at 80 rpm. On day 5, the properly developed Ebs were selected and transferred into an adherent 6-well plate with differentiation medium (X-vivo; Lonza) supplemented with 1% penicillin/streptomycin, 1 mM L-glutamine, 0.05 mM β-mercaptoethanol, 50 ng/mL M-CSF (Peprotech), and 25 ng/mL IL-3 (Peprotech). In case of feeder-free iPSC culture conditions, the protocol was modified accordingly [4,5] and to promote hematopoietic differentiation the mesoderm priming medium was supplemented with the cytokines BMP4, SCF and VEGF. The medium of the differentiation cultures was replaced once weekly and the generated macrophages were harvested and terminally differentiated in RMPI medium supplemented with 10 % FCS (Sigma Aldrich), 1 % penicillin/streptomycin and 50 ng/mL M-CSF (Peprotech) (0.1 x $10^6$ cells/well a 48-well plate).

**SDS-PAGE and western blot**

Protein samples were separated by SDS-PAGE in gels containing 1% TCE[62]. After the run, gels were transferred into VE-water and imaged after 3× 45 s activation using the stain-free gel settings at a ChemiDoc™ MP Imaging System. Alternatively, Coomassie staining was performed to visualize total protein. The proteins were then transferred onto nitrocellulose membranes (PALL via VWR #66485) and imaged for total protein in VE-water using the ChemiDoc stain-free settings, and then blocked with 5% milk in PBS-T (0.1% Tween20) for 1 h at RT. Subsequently, primary antibodies (anti-ICAM1, Cell Signalling Technologies #4915S and anti-β-actin, Thermo Scientific #MA 1-140) in 5% (w/v) milk in DPBS-T were added and incubated overnight at 4 °C. After washing 4 times, the membrane was incubated with secondary antibodies (Goat αMurine IgG IRDye680RD, LI-COR #925-68070 and Goat αRabbit IgG IRDye800CW, LI-COR #925-32211) in 2.5% (w/v) milk in DPBS-T for 1 h at RT in the dark. Finally, the membrane was washed again 3 times with DPBS-T and was transferred into 1× DPBS for imaging at the ChemiDoc™ MP. Quantification was performed using the band and lane tool of Image Lab™ 6.0.1 software.
